# Commensal bacteria inhibit viral infections via a tryptophan metabolite

**DOI:** 10.1101/2024.04.21.589969

**Authors:** Danting Jiang, Nicole Soo, Chin Yee Tan, Sedem Dankwa, Hsuan-Yuan Wang, Barbara S. Theriot, Keyi Xue, Demitrius Hill, Amir Ardeshir, James R. Bain, Sarah M. Heston, Jillian H. Hurst, Nazema Y. Siddiqui, Koen K. A. Van Rompay, Luisa Cervantes-Barragan, Jonathan P. Smith, Rinn Song, Jatin Roper, Matthew S. Kelly, Michael G. Hudgens, Kristina De Paris, Sallie R. Permar, Ria Goswami, Neeraj K. Surana

## Abstract

Clinical outcomes following viral exposures exhibit substantial interindividual variability. Although developing evidence suggests commensal bacteria modulate viral infections, the specific bacteria and mechanisms remain underexplored. Here we define a pathway by which viral infections are inhibited by specific tryptophan-catabolizing bacteria. Using HIV as a model, we bioinformatically associated and experimentally validated several bacterial species that inhibited viral replication. This activity required the aromatic amino acid aminotransferase (ArAT) to metabolize tryptophan into 3-indolelactic acid, which agonizes the aryl hydrocarbon receptor (AhR). Given that AhR regulates multiple viral infections, we found that commensal bacteria also inhibit cytomegalovirus (CMV) in an ArAT-dependent manner. Finally, we confirmed that ArAT is associated with protection against disease outcomes in three distinct human cohorts at-risk for HIV, CMV, or symptomatic COVID-19. Taken together, our results provide mechanistic insight into how commensal bacteria impact viral infections, thereby adding to an emerging field focused on host–commensal–virus interactions.

## INTRODUCTION

Despite recent advances in vaccines and antiviral therapeutics, viral infections pose a continued threat to humanity^1–3^. For all viruses, there exists significant heterogeneity in clinical outcomes of exposed individuals. While some people remain uninfected, others have disease severities that span the entire spectrum from asymptomatic to fatal^4,5^. Although numerous host characteristics (e.g., age, genetics, immune status) and viral features (e.g., virulence factors, route and dose of infection) contribute to this vast interindividual variability in clinical outcomes^4,6^, most of these aspects are difficult to meaningfully alter. Identifying modifiable factors that skew the response towards increased resistance to infection and/or severe disease, particularly if effective against multiple classes of viruses, may yield novel broadly-acting antiviral therapeutics^7^.

It is now clear that environmental factors, including the microbiota, also modulate susceptibility to viral infections^8–10^. Commensal bacteria can potentiate viral infectivity by the virus binding to bacterial surface polysaccharides, which help stabilize the virion^11–13^. Moreover, commensal bacteria play a fundamental role in setting the immunological tone and regulating immune responses^14,15^, which provides another mechanism for influencing the severity of viral disease, especially through modulation of the interferon pathway^13,16–20^. However, the bacterial mechanisms underlying these intricate microbiota–host–virus interactions remain poorly defined^8,9^. Identification of specific commensal bacteria-derived products that modify cellular susceptibility to viral infection could help explain the disparity in clinical outcomes of individuals with similar risk factors and exposures.

In earlier work, we noted that infant rhesus macaques varied greatly in their susceptibility to oral infection with simian–human immunodeficiency virus (SHIV)^21^, a model of breast milk transmission of HIV. In the present study, we correlated their susceptibility to SHIV acquisition with their fecal microbiota. We found that increased resistance to infection was bioinformatically associated with the presence of two bacterial taxa, *Lactobacillus gasseri* and the family Lachnospiraceae, which we then experimentally validated as inhibiting HIV infection in vitro and ex vivo. Ultimately, we delineated a pathway—including the bacterial gene, bacterial metabolite, and host receptor—by which specific commensal bacteria inhibit replication by different classes of viruses. Importantly, we confirmed the key aspect of our findings in four different human cohorts of individuals at-risk for infection with HIV, CMV, or SARS-CoV-2, thus generalizing our results to humans. Taken together, our study provides mechanistic insights into how the microbiome drives heterogeneous clinical outcomes resulting from viral exposures.

## RESULTS

### Microbe–phenotype triangulation associated two bacterial taxa with protection against SHIV infection

To identify commensal bacteria that protect against viral infections, we leveraged samples from a previously conducted study where infant rhesus macaques were orally challenged with SHIV to mimic breast milk transmission of HIV^21^. In brief, newborn animals were immunized with 3 doses of either a placebo or HIV envelope glycoprotein 120 (gp120)-based vaccine (*n*=12 animals/group) and, beginning at 15 weeks of age (3 weeks after the final immunization), received weekly oral challenges with SHIV until they became infected **(Figure 1A)**. Although the HIV vaccine did not confer protection against infection, the animals exhibited drastic variability in the time to acquisition of SHIV **(Figure 1B)**^21^. Since gastrointestinal tissues are one of the first sites that get infected with orally acquired virus^22^, we reasoned that differences in the intestinal microbiome may contribute to the variability in time to acquisition of SHIV.

**Figure 1.**
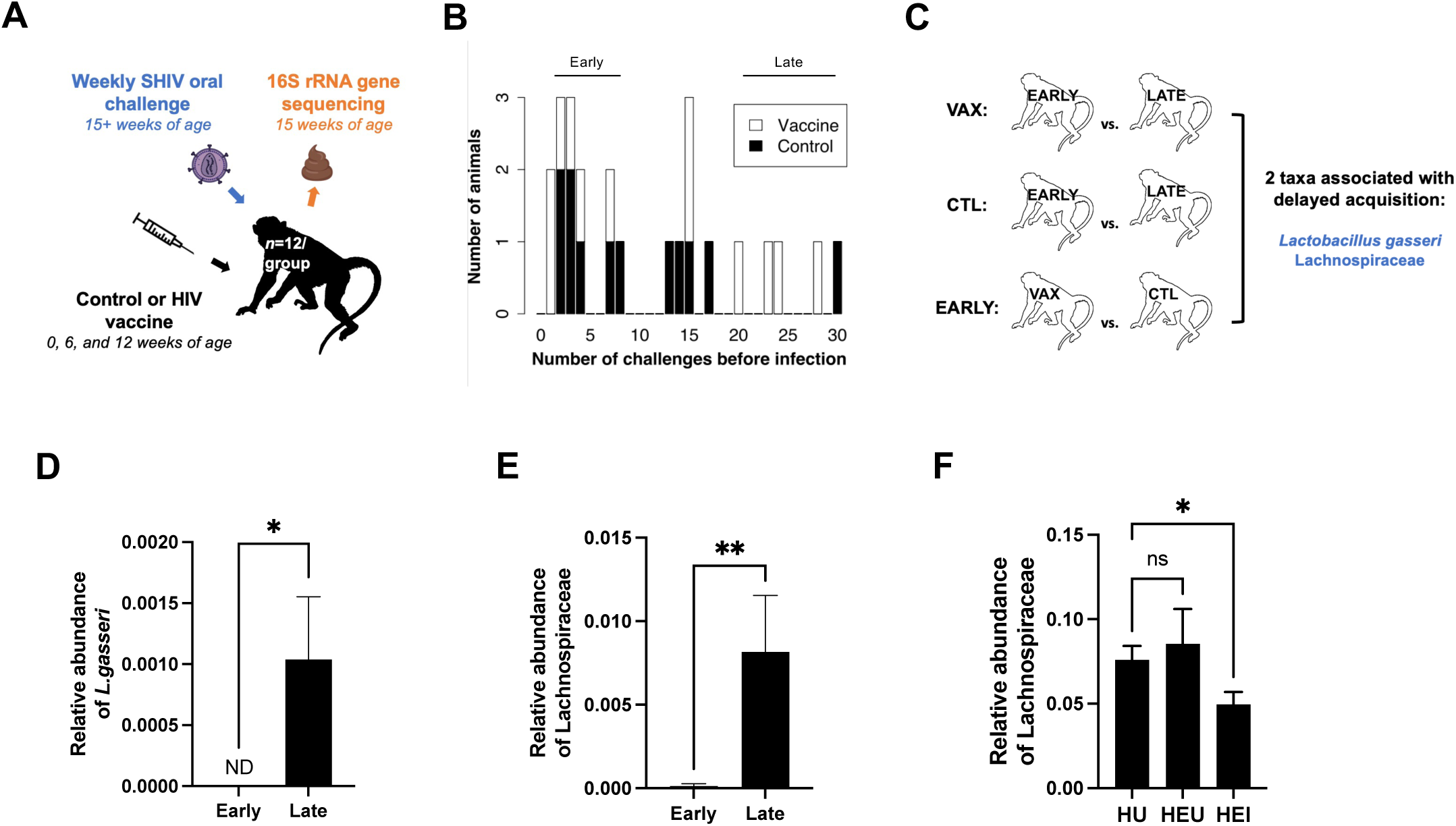
Microbe–phenotype triangulation associated two bacterial taxa with protection against SHIV infection. **(A)** Schematic of the nonhuman primate study design. Newborn animals were immunized by either an HIV or control vaccine (*n*=12 animals/group) at 0, 6, 12 weeks of age. Starting at 15 weeks of age, animals were orally challenged with SHIV weekly until they became infected. Fecal samples were collected just prior to the SHIV challenges for 16S rRNA gene sequencing. **(B)** Distribution of the number of weekly SHIV challenges each monkey received before becoming infected. Bars are colored by regimen groups. Data is from reference 21. **(C)** The different pairwise comparisons used for microbe-phenotype triangulation are depicted. The two bacterial taxa associated with delayed acquisition of SHIV are listed. VAX, animals receiving the vaccine against HIV; CTL, animals receiving the control vaccine. **(D–E)** Relative abundance of *L. gasseri* **(D)** and Lachnospiraceae **(E)** in HIV-vaccinated animals that were infected early (≤8 challenges) or late (≥20 challenges). **(F)** Relative abundance of the family Lachnospiraceae in the fecal microbiota of infants and very young children with different HIV statuses. HU, HIV unexposed (*n*=91); HEU, HIV-exposed uninfected (*n*=16); HEI, HIV-exposed infected (*n*=28). The bars in panels D–F represent means±SEM. ND, not detected; ns, not significant; \**P*<0.05; \*\**P*<0.01 by Mann-Whitney U test **(D–E)** or by Welch’s ANOVA with post-hoc Dunnett’s tests **(F)**.

Accordingly, we analyzed the fecal microbiome of these rhesus macaques at 15 weeks of age— just before they began the oral SHIV challenge—to identify bacterial taxa that are associated with delayed acquisition of SHIV. The time to SHIV acquisition in these animals had a wide distribution of 1–30 challenges (**Figure 1B)**. We reasoned taxa that are differentially abundant in animals at the two extremes of this spectrum—i.e., those infected early (≤8 challenges) or late (≥20 challenges)—may be related to differences in viral susceptibility. We assessed the vaccination and control groups separately since we previously demonstrated that vaccination altered the microbiota in these animals^23^. For the control group of animals, there was only 1 animal in the late-infected group, which precluded our ability to identify differentially abundant taxa. By comparing the microbiota in the HIV-vaccinated animals that got infected early with the HIV-vaccinated animals that got infected late, we identified 4 taxa that were more abundant in the late-infected animals **(Supplementary Table 1).** Applying concepts of microbe–phenotype triangulation^24^, we eliminated 2 of these taxa that were differentially abundant in the microbiomes of early-infected animals in the HIV vaccine and control groups as the abundance of these taxa are driven by the effect of vaccination without impacting susceptibility to infection (**Figure 1C; Supplementary Table 1**). Ultimately, we identified two taxa—*Lactobacillus gasseri* and the family Lachnospiraceae—that were bioinformatically associated with delayed acquisition of SHIV **(Figure 1C–E)**. Notably, *L. gasseri* has previously been identified as inhibiting HIV infection in vitro^25,26^, which supports the idea that our bioinformatic analyses identified relevant taxa.

### Lachnospiraceae is associated with protection against HIV infection in humans

To determine whether our findings may be relevant to humans, we sought to identify a population that is equally at-risk for acquiring HIV but has discordant outcomes such that we can determine whether *L. gasseri* or Lachnospiraceae is more abundant in those who do not acquire HIV. We reasoned that children born to mothers with HIV represent a group that would facilitate this analysis. As such, we analyzed the fecal microbiota of Kenyan infants and very young children exposed to HIV perinatally who either did (*n*=28) or did not acquire HIV (*n*=16)^27^, with Kenyan children not exposed to HIV serving as a control (*n*=91). Although *L. gasseri* was not detected in any of the samples (potentially due to the young age of the cohort), we found that HIV-unexposed (HU) and HIV-exposed uninfected (HEU) children had a greater fecal abundance of Lachnospiraceae than those children who perinatally acquired HIV (HEI; **Figure 1F**). These data intriguingly suggest that the bacterial family Lachnospiraceae is associated with protection against HIV infection in humans, similar to our findings in infant rhesus macaques.

### Human-derived *L. gasseri* and Lachnospiraceae isolates robustly inhibit HIV infection

We obtained human-derived isolates of the identified bacterial taxa to experimentally validate whether these bioinformatic associations also represent causal relationships. We treated TZM-bl cells with heat-killed cultures of these bacteria for 48 hours before infecting with HIV to assess their impact on viral replication. TZM-bl cells are sensitive to infection with diverse HIV-1 isolates due to their high-level expression of CD4, CXCR4, and CCR5^28^. Moreover, these cells express luciferase under control of the HIV-1 long terminal repeat, which permits sensitive and accurate measurements of infection^29^. To control for the potential of non-specific effects due to the addition of bacteria, we normalized the effects of our bacteria of interest to TZM-bl cells treated with *Bacteroides fragilis*, a bacterium commonly found in the intestinal microbiota and used here as a negative control; none of the bacteria tested induced significant cytotoxicity **(Figure S1A)**. While *L. gasseri* exhibited strain-dependent inhibition against HIV infection, two Lachnospiraceae isolates, *Clostridium immunis* and *Ruminococcus gnavus*, robustly inhibited HIV replication irrespective of virus tropism or viral clade **(Figure 2A–B** and **Figure S1B–D)**, with levels of inhibition approaching that of the antiretroviral (ARV) control. Importantly, *C. immunis* and *R. gnavus* also inhibited HIV infection of primary CD4+ T cells present in tonsillar mononuclear cells (TMCs) isolated from HIV-naïve children (**Figure 2C**). Given that *L. gasseri* had variable efficacies in these in vitro and ex vivo assays, we focused on *C. immunis* and *R. gnavus* in subsequent experiments.

**Figure 2.**
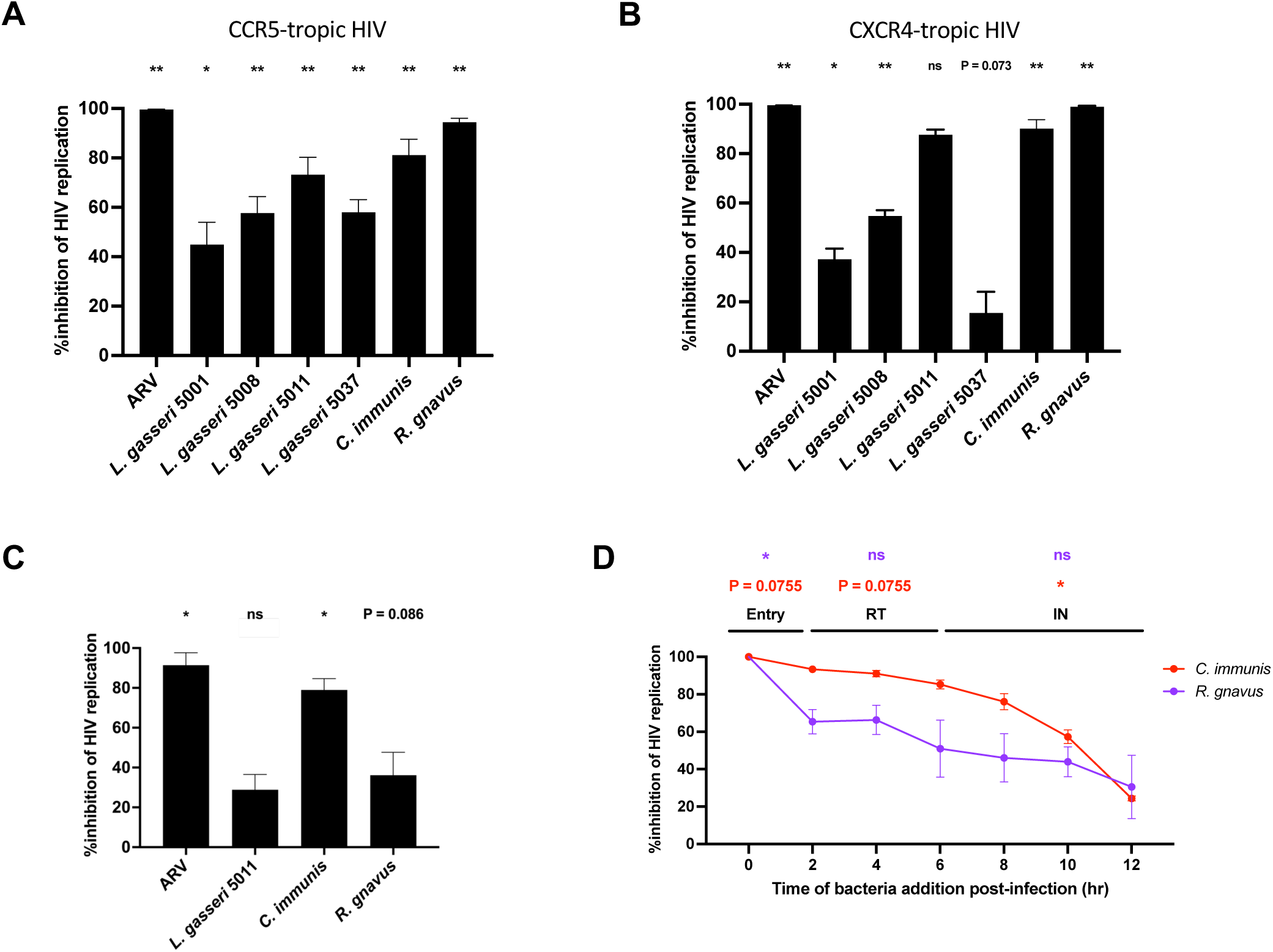
Human-derived *L. gasseri* and Lachnospiraceae isolates robustly inhibit HIV infection. **(A–B)** Percent inhibition of HIV replication for CCR5-tropic **(A)** and CXCR4-tropic **(B)** strains of HIV in TZM-bl cells treated with the indicated commensal bacteria. ARV, antiretroviral cocktail. Data are normalized to HIV-infected TZM-bl cells treated with *B. fragilis*. **(C)** Percent inhibition of HIV replication in human tonsillar CD4+ T cells treated with the indicated commensal bacteria. Data are normalized to tonsillar CD4^+^ T cells treated with *B. fragilis*. **(D)** Time-of-addition assay results for *C. immunis* and *R. gnavus*. The times depicted for the various viral pathogenesis steps were determined using antiviral drugs targeting each step. RT, reverse transcription; IN, integration. The data for each curve is normalized to TZM-bl cells treated with the bacteria at 12 hours after infection with HIV. Panels A–D include data pooled from 6, 6, 4, and 3 independent experiments, respectively; in panel B, the data for *L. gasseri* 5011 is from 3 independent experiments. Data are represented as mean±SEM. *P*-values between 0.05 and 0.1 are indicated explicitly on each figure. ns, not significant; \**P*<0.05; \*\**P*<0.01 by paired t-tests for each treatment compared to the negative control **(A–C)** or comparing the same bacterium added at specific timepoints (0 vs. 2 hr for entry, 2 vs. 6 hr for RT, and 6 vs. 12 hr for IN; **D**). *P* values were adjusted for multiple comparisons by the Benjamini-Hochberg procedure. See also **Figures S1 and S2**.

To gain insight into the mechanism of action of these commensal bacteria, we performed time-of-addition (TOA) assays to define which step(s) of HIV pathogenesis is impacted^30^. This TOA approach assesses how long the addition of an antiviral agent can be delayed without losing its efficacy. We used several antiretroviral drugs with known mechanisms of action to determine the post-infection time points that reflect the viral replication steps they target (viral entry/fusion with the host cell, reverse transcription, and integration; **Figure S2)**. Comparing the TOA results for the bacteria with that of the controls, we found that *R. gnavus* significantly inhibits the viral entry/fusion step. In contrast, *C. immunis* exhibited a steady loss of inhibitory activity throughout 12 hours post-infection (**Figure 2D**), which demonstrates an impact on all three steps. Although the largest decrease in activity is during the integration phase, this may simply be a reflection of how these TOA assays work, with less pronounced effects observed in early steps when multiple steps are impacted^30,31^. Given that *C. immunis* robustly inhibits HIV in vitro and ex vivo and impacts multiple steps of HIV replication, we focused on this bacterium as we sought to determine the basis of commensal bacteria-mediated antiviral effects.

### Commensal bacteria inhibit HIV infection via the ArAT

It has long been appreciated that the degree of tryptophan metabolism is linked to HIV pathogenesis and disease severity, with increases in the serum kynurenine:tryptophan (K/T) ratio linked to progressive disease^32,33^. These studies have largely focused on host-mediated catabolism of tryptophan to kynurenine by indoleamine-2,3-dioxygenase (IDO1); however, the gut microbiota can also breakdown tryptophan into different metabolites through the bacterial aromatic amino acid aminotransferase (ArAT)^34^. Along these lines, *C. immunis* has an ArAT gene, which we hypothesized is critical for its ability to inhibit HIV infection by regulating tryptophan metabolism. If true, we speculated that the active component of *C. immunis* should be present in the bacterial supernatant. Indeed, while the bacterial supernatant was able to inhibit HIV infection of TZM-bl cells to a similar extent as the unfractionated culture, the *C. immunis* cell pellet lacked this antiviral activity (**Figure 3A**). These data indicate that *C. immunis* secretes some factor into the supernatant that prevents HIV infection.

**Figure 3.**
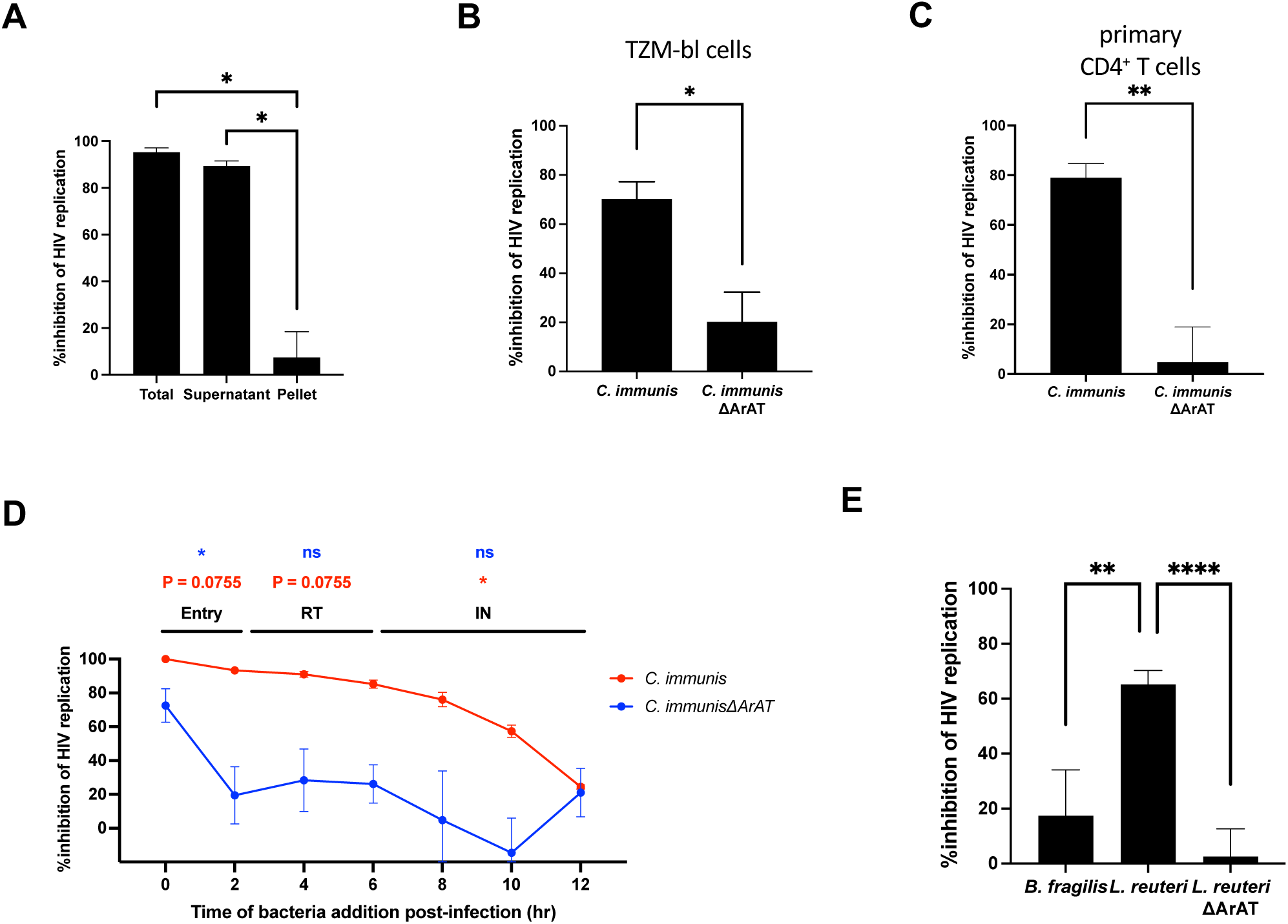
Commensal bacteria inhibit HIV infection via the ArAT. **(A)** Percent inhibition of HIV replication in TZM-bl cells treated with the indicated fraction of heat-killed *C. immunis* culture. Total refers to unfractionated culture. **(B–C)** Percent inhibition of HIV replication in TZM-bl cells **(B)** or human tonsillar CD4^+^ T cells **(C)** treated with *C. immunis* or *C. immunis*ΔArAT. The data for *C. immunis* in panel C are the same as that presented in Figure 2C. **(D)** Time-of-addition assay results for *C. immunis* and *C. immunis*ΔArAT. The times depicted for the various viral pathogenesis steps were determined using antivirals targeting each step. RT, reverse transcription; IN, integration. The data for *C. immunis* are the same as that presented in Figure 2D. **(E)** Percent inhibition of HIV replication in TZM-bl cells treated with *B. fragilis*, *L. reuteri*, or *L. reuteri*ΔArAT. Data are normalized to HIV-infected TZM-bl cells or primary CD4^+^ T cells treated with *B. fragilis* **(A–C)**, TZM-bl cells treated with the indicated bacteria at 12 hours post-infection **(D)**, or HIV-infected TZM-bl cells not treated with bacteria **(E)**. In panels A–E, data are pooled from 5, 4, 4, 3, and 7 independent experiments, respectively. Data are represented as mean±SEM. *P*-values between 0.05 and 0.1 are indicated explicitly on each figure. ns, not significant; \**P*<0.05; \*\**P*<0.01; \*\*\*\**P*<0.0001 by paired t-tests for the displayed comparisons **(A–C, E)** or comparing the same bacterium added at specific timepoints (0 vs. 2 hr for entry, 2 vs. 6 hr for RT, and 6 vs. 12 hr for IN; **D**). *P* values in panels A, D, and E were adjusted for multiple comparisons by the Benjamini-Hochberg procedure. See also **Figure S2**.

To more directly test the role of *C. immunis* ArAT in HIV pathogenesis, we sought to knockout the gene. Although genetic manipulation of most commensal bacteria remains challenging^35,36^, we developed new genetic tools for *C. immunis* and generated an isogenic mutant that lacks a functional ArAT gene (*C. immunis*ΔArAT). Remarkably, the *C. immunis*ΔArAT mutant had greatly reduced ability to inhibit HIV infection in TZM-bl cells or in primary CD4+ T cells present in human TMCs compared to wildtype *C. immunis* **(Figure 3B–C)**. TOA studies performed with the *C. immunis*ΔArAT mutant demonstrated that it inhibits viral entry/fusion but not any post-fusion steps (**Figure 3D**), a finding that indicates ArAT is involved in limiting these later steps of HIV pathogenesis.

Given that the ArAT gene is present in diverse bacteria, we reasoned that other commensal bacteria may also inhibit HIV infection in an ArAT-dependent manner. Accordingly, we tested whether *Lactobacillus reuteri* and an isogenic mutant deficient in ArAT (*L. reuteri*ΔArAT) are able to inhibit HIV infection. Intriguingly, while *L. reuteri* possessed inhibitory activity, *L. reuteri*ΔArAT did not **(Figure 3E)**. Taken together, our data establish that taxonomically disparate commensal bacteria can inhibit HIV replication via ArAT.

### Commensal-derived indole lactic acid inhibits HIV infection by activating the aryl hydrocarbon receptor

We performed untargeted metabolomics on the culture supernatant from *C. immunis* and *C. immunis*ΔArAT to identify compounds that might explain the difference in activity. Of the 214 annotated metabolites, only 1—indole-3-lactic acid (ILA)—was differentially abundant between the two strains, with the mutant unable to produce any ILA (**Figure 4A**). Of note, *L. reuteri* has also been shown to produce ILA, in addition to indole-3-aldehyde (I3A), in an ArAT-dependent manner^37,38^. We treated cells with each of these compounds to determine whether either is sufficient for inhibiting HIV infection. Interestingly, we found that ILA—but not I3A—was sufficient to inhibit HIV infection **(Figure 4B)**, a finding that suggests there is some specificity in terms of which tryptophan catabolites have antiviral activity. Given that ILA is a known agonist of the aryl hydrocarbon receptor (AhR)^34^, a transcription factor that regulates expression of a broad array of genes, we hypothesized that bacterial ArAT-derived ILA inhibits HIV infection by activating this pathway. Indeed, the canonical AhR agonist 6-formylindolo(3,2-b)carbazole (FICZ) inhibits HIV infection while the AhR antagonist CH223191 enhanced HIV infection **(Figure 4B)**. These findings highlight the importance of AhR activity in regulating HIV infection.

**Figure 4.**
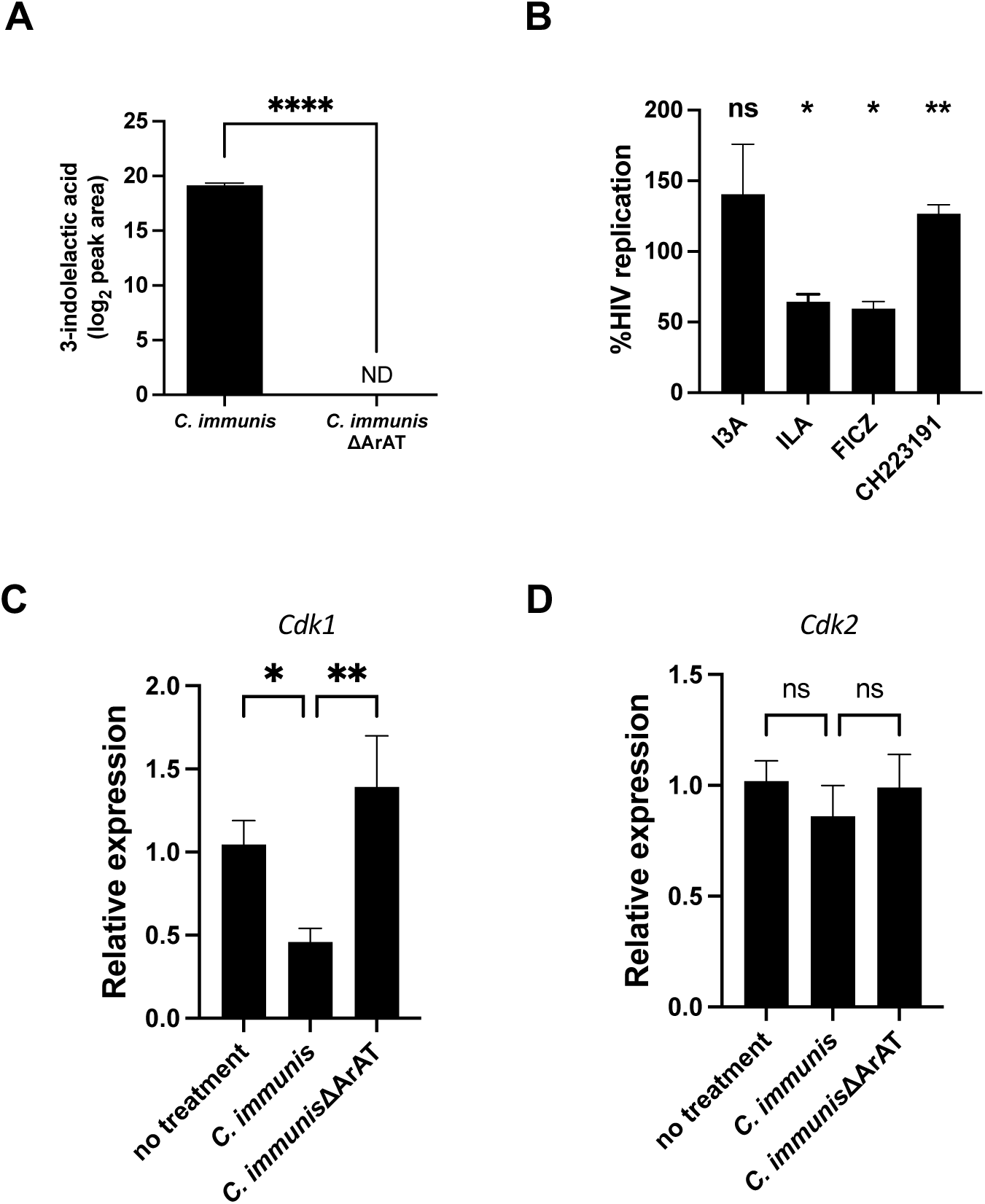
Commensal-derived indole lactic acid inhibits HIV infection by activating the aryl hydrocarbon receptor. **(A)** Mass spectrometric quantitation of ILA produced by *C. immunis* and *C. immunis*ΔArAT. **(B)** Percent HIV replication in TZM-bl cells treated with the indicated small molecule. Data are normalized to cells treated with DMSO. **(C–D)** Relative expression of *Cdk1* **(C)** and *Cdk*2 **(D)** in HIV-infected TZM-bl cells treated with the indicated bacteria. Data are normalized to HIV-infected cells without bacterial treatment. In panels A, C, and D, data are from 7, 6, and 6 independent experiments, respectively; in panel B, data are from 3 independent experiments for I3A, 5 for ILA, 4 for FICZ, and 11 for CH223191. Data are represented as mean±SEM. ND, not detected; ns, not significant; *, *P*<0.05; **, *P*<0.01; ****, *P*<0.0001 by two-sample t-test **(A)**, paired t-tests for each treatment compared to the negative control and adjusted by the Benjamini-Hochberg procedure **(B)**, or Kruskal-Wallis with post-hoc Dunn’s tests **(C–D)**.

Importantly, activation of the AhR reduces viral infections by inhibiting expression of cyclin-dependent kinases (*Cdk*) 1 and 2 along with their associated cyclins^39^. Consistent with this, we found that *C. immunis* treatment of TZM-bl cells led to decreased expression of *Cdk1* with no impact on *Cdk2* **(Figure 4C–D)**. Moreover, this effect on *Cdk1* expression was dependent on ArAT **(Figure 4C)**. Collectively, these data demonstrate that commensal bacteria-derived ILA activates the AhR, which leads to decreased expression of *Cdk1* and inhibition of HIV replication.

### The abundance of the ArAT gene is lower in humans who develop viral infections

To determine whether our findings may be relevant to human infections, we analyzed publicly available fecal shotgun metagenomic data from a longitudinal study that compared the microbiome in 9 men who have sex with men (MSM) before and immediately after HIV seroconversion with that of 20 individuals from a similar high-incidence MSM population who did not acquire HIV^40^. For the individuals who acquired HIV, we focused our analysis on the samples obtained at the visit immediately preceding their documented seroconversion. Consistent with our earlier findings **(Figure 1F)**, individuals who acquired HIV had a lower abundance of Lachnospiraceae prior to HIV seroconversion compared to those who remained HIV-negative throughout the study **(Figure S3A)**. Intriguingly, those individuals who ultimately acquired HIV also had a lower abundance of the ArAT gene compared to those who did not acquire HIV **(Figure 5A)**, with a trend towards lower serum ILA levels **(Figure S3B)**. Taken together, these findings suggest the bacterial ArAT gene is associated with protection against HIV in humans.

**Figure 5.**
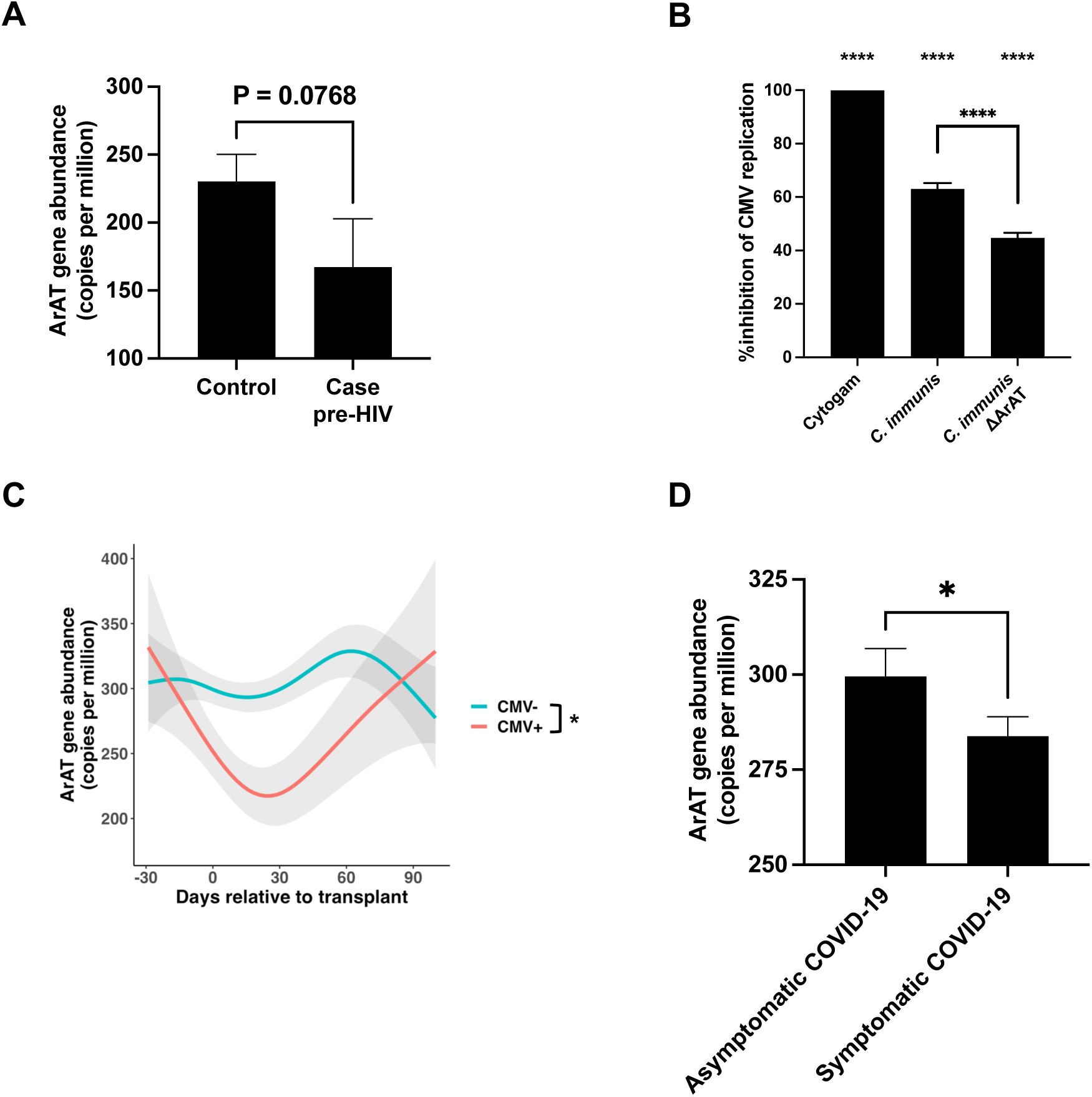
The abundance of the ArAT gene is lower in humans who develop viral infections. **(A)** The fecal abundance of the ArAT gene in adults at-risk for HIV who did (case; *n* = 9) or did not acquire HIV (control; *n* = 20). The samples for the cases were obtained before they tested positive for HIV. **(B)** Percent inhibition of CMV replication in ARPE-19 cells treated with CMV immunoglobulin (Cytogam), *C. immunis*, or *C. immunis*ΔArAT. Data are normalized to ARPE-19 cells treated with *B. fragilis*. Data are from 7 independent experiments, with bars representing means±SEM. **(C)** The abundance of the ArAT gene in longitudinal fecal samples of children undergoing hematopoietic stem cell transplantation who do (*n* = 13) or do not develop detectable CMV (*n* = 59). Lines with shaded areas represent means±SEM calculated from a generalized additive mixed model with fixed effect for CMV status, random effect for each individual, and a smooth term for days relative to transplant. **(D)** The fecal abundance of the ArAT gene in individuals infected with SARS-CoV-2 who did (*n* = 166) or did not develop COVID-19-related symptoms (*n* = 67). *, *P*<0.05; ****, *P*<0.0001 by a Mann-Whitney U test **(A and D)**, paired t-tests **(B)**, or a Wald test for CMV status **(C)**. In panel B, the asterisks at the top of the graph represent the *P* value obtained when comparing each sample to the control by paired t-tests and adjusted by the Benjamini-Hochberg procedure; the asterisks above the bracket represent the *P* value obtained when comparing *C. immunis* and *C. immunis*ΔArAT by a paired t-test. See also **Figures S3 and S4**.

We speculated this commensal bacteria–ILA–AhR pathway may be relevant in modulating the susceptibility to other viruses given that activation of the AhR has been linked to the pathogenesis of numerous viral infections (e.g., human cytomegalovirus [HCMV], Zika virus, SARS-CoV-2)^41,42^. Accordingly, we tested whether *C. immunis* can modulate HCMV infection in vitro. Remarkably, *C. immunis* inhibited HCMV infection in an ArAT-dependent manner **(Figure 5B)**, a finding that demonstrates bacterial ArAT can inhibit infection from vastly divergent viruses. To determine whether the abundance of the ArAT gene is differentially abundant in individuals with or without CMV, we analyzed fecal shotgun metagenomic data generated from 72 children undergoing hematopoietic stem cell transplantation^43^, an immunosuppressed population who are high-risk for CMV in the time period immediately following their transplant^44^. Longitudinal stool samples were acquired weekly beginning up to one month prior to transplant and continued until 3 months after transplant. Although all children had a similar abundance of the ArAT gene preceding their transplant, its abundance immediately after transplant decreased specifically in children who developed detectable CMV viremia **(Figure 5C)**, a finding that suggests the decrease in ArAT gene abundance may be associated with CMV viremia. Interestingly, the abundance of the ArAT gene in the CMV viremic cohort normalized by 2–3 months after transplant, a time at which their risk for CMV generally decreases.

Given these findings for HIV and CMV, we wondered whether bacterial ArAT may also play a role in other viral infections. Accordingly, we compared fecal shotgun metagenomic data from 285 individuals who were prospectively enrolled after a household exposure to SARS-CoV-2^45^, another virus known to be impacted by the AhR signaling pathway^42^. In contrast to our observations with HIV and CMV, we found no difference in the fecal abundance of the ArAT gene between participants who did or did not develop infection with SARS-CoV-2 **(Figure S4)**. However, given previous work demonstrating that COVID-19 disease severity is linked to changes in the gut microbiome^46,47^, we reasoned that differences in fecal ArAT gene abundance may be associated with the severity of COVID-19. We stratified individuals by the presence or absence of symptoms as all the SARS-CoV-2-infected participants in our cohort were either asymptomatic or had mild disease^45^. Interestingly, we found that the abundance of the ArAT gene was significantly greater in people with asymptomatic COVID-19 compared to those who became symptomatic **(Figure 5D)**. Taken together, these findings strongly support the notion that bacterial ArAT-mediated metabolism of tryptophan protects against viral diseases.

## DISCUSSION

Starting with infant rhesus macaques that exhibited differential susceptibility to enteral SHIV infection^21^, we bioinformatically identified and experimentally validated several bacterial species as inhibiting HIV infection in vitro and ex vivo. We further delineated a pathway where specific commensal bacteria use the ArAT to catabolize tryptophan into ILA that activates the AhR and results in increased resistance against viral infections. Importantly, we leveraged metagenomic data from 4 different cohorts where individuals were at-risk for HIV, CMV, or SARS-CoV-2 to demonstrate our findings are generalizable to humans.

Our study presents a highly effective framework to identify specific bacteria and the underlying molecular mechanisms that are causally related to a phenotype of interest, something that remains challenging to accomplish in microbiome studies^48,49^. Our successful implementation of microbe-phenotype triangulation, an innovative bioinformatic discovery platform we previously developed^24^, to associate two bacterial taxa with increased resistance to viral infection further demonstrates the utility of this approach^24,50^. Indeed, our analysis independently identified *L. gasseri*, which has previously been shown to inhibit HIV infection in vitro^25,26^. Moreover, our association of Lachnospiraceae with increased resistance to viral infection is consistent with its abundance being lower in individuals living with HIV than in healthy controls^51–55^, which we additionally confirmed in perinatally exposed children and an MSM population who developed discordant HIV statuses. Similar to our previous work^24^, we were able to increase the specificity of our bioinformatic analyses by incorporating pairwise comparisons that should not relate to the phenotype of interest. Here, we were able to exclude two *Bacteroides* spp. from consideration, and we ultimately used a *Bacteroides* sp. as our experimental negative control to further demonstrate that our taxa of interest had biologic effects that are not present in all bacteria.

We extended our discovery pipeline by combining it with principles of microbial pathogenesis to elucidate the underlying molecular mechanism. We developed bacterial genetics for *C. immunis*, a human-derived Lachnospiraceae isolate, so we could generate an isogenic mutant lacking ArAT, which catalyzes the first step of tryptophan catabolism to specific indolic compounds^34^. Interestingly, the *C. immunis*ΔArAT mutant had decreased inhibitory activity against HIV and CMV infections compared to wild-type *C. immunis*. While this type of genetic inactivation is still infrequent in microbiome studies^48^, it will become more commonplace as genetic tools for manipulating commensal bacteria become more advanced^56,57^. Identifying a specific gene as being critical to inhibiting viral infections enabled us to investigate the abundance of the ArAT gene in shotgun metagenomic data derived from relevant patient populations to evaluate relevance in human disease. Although we analyzed fecal metagenomic data for each of our cohorts, analysis of samples from disease-relevant sites (e.g., respiratory samples for patients with respiratory tract infections) may provide greater sensitivity. Taken together, we have combined microbe–phenotype triangulation as a discovery platform, bacterial genetics, and metagenomic analyses to elucidate the mechanism and translational relevance of microbiome-mediated inhibition of viral infections.

We found that the *C. immunis* ArAT produced ILA, a microbiome-derived endogenous agonist for the AhR signaling pathway, that protects against viral infections. The AhR is strongly linked to viral pathogenesis^42^, though there is ongoing controversy whether the AhR pathway promotes or restrains these infections^41,42^. These discrepancies may relate to the role of AhR signaling varying in a virus-^42,58–61^, cell type-^58,62,63^, and ligand-specific manner^62,64,65^. Consistent with these findings, we found that ILA—but not I3A—robustly suppresses viral infection even though both are AhR agonists that are produced in an ArAT-dependent manner^34,37^. It is likely that other microbiome-derived AhR agonists may similarly inhibit viral infections, though additional studies are needed to identify the specific subset of AhR agonists with this activity.

In addition to microbiome-mediated tryptophan catabolism, the host can also metabolize tryptophan directly, with generation of kynurenine by IDO1 being a common pathway. The level of kynurenine, often reported as the ratio of kynurenine to tryptophan (K/T ratio), is linked to more severe viral infections^32,33,66,67^. Although kynurenine does not have direct AhR binding activity, it is an AhR pro-ligand that requires chemical conversions to act as an AhR agonist^68^. While kynurenine can modulate the immune system in an AhR-dependent manner^69–71^, it is less clear whether it directly impacts the severity of viral infections in vivo^72–74^. We speculate that the clinical observations linking kynurenine to more severe viral disease may be driven by differences in microbiome-mediated tryptophan catabolism, with increased kynurenine levels serving as a proxy for lower levels of tryptophan-catabolizing bacteria. Metabolism of tryptophan by commensal bacteria, which occurs in the intestinal lumen prior to absorption of tryptophan, may reduce the amount of tryptophan available to the host for conversion into kynurenine.

It is notable that commensal bacteria-derived ILA can inhibit both HIV, a retrovirus, and CMV, a DNA herpesvirus, given the vast differences in their underlying biology. These effects do not require modulation of the immune system as the effects are also present in epithelial cell line-based experiments, a finding that suggests the bacteria are altering cellular physiology in a way that affects viral pathogenesis. AhR activation is known to modulate expression of numerous genes, including repressing expression of *Cdk1* and *Cdk2*^39^. Along these lines, we found that cells treated with *C. immunis* have reduced expression of *Cdk1* in an ArAT-dependent manner, a finding that helps mechanistically explain how commensal bacteria-generated ILA leads to protection against viral infection as inhibition of CDK1 has been shown to reduce viral replication^75,76^. CDK1 normally phosphorylates SAMHD1^77^, thereby inhibiting its dNTPase activity^77,78^; decreased *Cdk1* expression leads to an increase in active, non-phosphorylated SAMHD1 and a concomitant reduction in the dNTP pool available for viral DNA synthesis^79,80^. This mechanism, which is similar to what has been previously demonstrated for FICZ^39^, is consistent with our finding that the *C. immunis* ArAT impacts the reverse transcription step of the HIV life cycle. We speculate that *C. immunis*, *L. reuteri*, and other ArAT-expressing commensal bacteria inhibit viral infection for a broad array of viruses—beyond just HIV and CMV—since the underlying mechanism revolves around the availability of dNTPs necessary for viral replication.

Overall, by starting with the observation that infant rhesus macaques had variable susceptibility to SHIV challenges, we defined a mechanistic pathway for how the microbiota influences the outcome of viral exposures. We used microbe–phenotype triangulation to associate bacterial taxa with enhanced resistance to viral infection, in vitro and ex vivo experiments to validate our bioinformatic findings, bacterial genetics to clarify the critical role of the ArAT gene in multiple bacteria, and metagenomic analyses of four distinct human cohorts to demonstrate the translational relevance of our findings. Taken together, our results provide a mechanistic basis for the significant heterogeneity in clinical outcomes of virus-exposed individuals. More broadly, our findings offer the potential for a microbiome-based therapeutic to treat and/or prevent viral infections.

## METHODS

### Microbe–phenotype triangulation to identify protective taxa in SHIV-challenged monkeys

We used data from a previous study involving a cohort of infant rhesus macaques that were challenged with simian–human immunodeficiency virus (SHIV) and exhibited variable levels of susceptibility to infection^21^. As previously described^21^, animals were immunized with either a HIV gp120-based vaccine or control vaccine at 0, 6, and 12 weeks of age (*n* = 12/group) before they were orally challenged with SHIV weekly beginning at 15 weeks of age. The fecal microbiome of these animals at 15 weeks of age has been previously described^23^. Briefly, they were generated by 16S rRNA gene sequencing, with data processed by the standard DADA2-based pipeline and taxonomy assignment based on the SILVA 132 database^81^. Identification of differentially abundant taxa for the pairwise comparisons used in the microbe–phenotype triangulation approach was accomplished in R (version 4.2.2) using DESeq2 with default thresholds^82^. Taxa present in <50% of animals in both groups of the comparison were excluded from the analysis.

### 16S rRNA gene sequencing analysis to detect Lachnospiraceae in Kenyan children

We obtained fecal samples from a previously reported cohort of Kenyan children (<5 years old) who had discordant HIV statuses^27^. DNA was extracted from the fecal samples using a MagAttract PowerSoil DNA EP Kit (Qiagen) according to the manufacturer’s instructions. The V4 region of the 16S rRNA gene was PCR-amplified using primers 515F and 806R as previously described^83^. Amplicons were quantified using a Qubit dsDNA HS assay kit (ThermoFisher), pooled in equimolar concentration, and sequenced on a MiSeq sequencer (Illumina; 2×250 base pair). Demultiplexed, adapter-free 16S rRNA gene sequencing data were imported into R (version 4.2.2) and processed by the standard pipeline of the DADA2 package^84^: paired-end reads were quality filtered, trimmed, denoised, and merged into an amplicon sequence variant (ASV) table followed by chimera removal and taxonomy assignment based on the SILVA 132 database^85^. Normalization of the ASV table by relative abundance was executed by the phyloseq R package^86^. We excluded samples from individuals who were less than 1-year-old and/or considered malnourished (i.e., weight-for-age z-score<-2) by the WHO Child Growth Standards (https://www.who.int/tools/child-growth-standards). We extracted the sum of the relative abundances of all ASVs classified within the family Lachnospiraceae. Use of Kenyan pediatric stool samples in this analysis was approved by the institutional review boards (IRBs) of the United States Centers for Disease Control and Prevention, the Kenya Medical Research Institute, and the Jaramogi Oginga Odinga Hospital. Written informed consent for the use of these stool samples was given by parents or legal guardians of participants.

### Bacterial strains

*C. immunis*, *R. gnavus* (ATCC 29149), and *B. fragilis* (NCTC 9343) were obtained from Dennis Kasper (Harvard University)^24^. All *L. gasseri* strains were isolated from catheterized urine samples of healthy women as previously described^87^; this study was approved by the Duke University Health System [DUHS] IRB (Pro00083917). *L. reuteri* 100-23 (wild-type and ΔArAT mutant) were previously described^37^. *C. immunis*, *R. gnavus*, and *B. fragilis* were grown in peptone–yeast– glucose (PYG) broth (Anaerobe Systems) or brain heart infusion-supplemented (BHI-S) broth (ATCC medium 1293). *Lactobacillus* strains were grown in De Man, Rogosa and Sharpe (MRS) broth (Sigma). All bacterial strains were grown overnight in an anaerobic chamber (Coy laboratories) with 2.5% H_2_ and 0 ppm O_2_ at 37°C. Serial dilutions of overnight cultures of the bacterial isolates were plated to enumerate the colony-forming units (CFU). Cultures were heat-killed at 65°C for 20 mins, aliquoted, and frozen at -80°C until further use. To separate the bacterial supernatant from the bacterial cells, the heat-killed cultures were centrifuged at 10,000 rpm for 5 min. The culture supernatant was collected, and the bacterial pellet was washed twice with PBS and resuspended in an equivalent volume of TZM-bl maintenance media.

### Cell lines and primary cells

TZM-bl luciferase reporter cells (BEI Resources) and human embryonic kidney-293T cells (HEK-293T; ATCC) were cultured in Dulbecco’s Modified Eagle Medium (DMEM; Gibco), with 10% heat-inactivated FBS (Millipore Sigma), 1% penicillin-streptomycin (Gibco), and 25 mM of HEPES (Gibco). Human retinal pigment epithelial cells (ARPE-19; ATCC) were cultured in DMEM-F12 (Gibco), with 10% FBS and 1% penicillin-streptomycin (Gibco). Human foreskin fibroblast cells (HFF-1; ATCC) were maintained in DMEM (Gibco) with 20% FBS, 1% penicillin-streptomycin, 2mM of L-glutamine, 25mM of HEPES, and 50 µg/ml of gentamicin. All cell lines and primary cells were cultured at 37°C in a humidified incubator with 5% CO_2_.

Tonsil tissues removed during routine tonsillectomy from HIV-naïve children (<15 years of age) were obtained through the Weill Cornell Medicine Department of Pathology and Laboratory Medicine. The use of tonsil specimens for this study was reviewed and determined to be exempt from human subject research by the Weill Cornell Medicine IRB; informed consent was therefore not required (Protocol # 21-08023851). Tonsil tissues were received within 1–3 hours of surgery; washed and minced in RPMI 1640 (Gibco) containing 1% antibiotic/antimycotic (Gibco), 50 μg/ml of gentamicin (Gibco), 10 μg/ml of cephalothin (Sigma-Aldrich), and 5 μg/ml of fluconazole (Sigma-Aldrich); and TMCs were isolated by filtering tonsillar tissue pieces through a 70 μm mesh filter, followed by density gradient centrifugation using Ficoll-Paque PLUS (GE Healthcare), as previously described^88^.

### Virus propagation and titration

The HIV-1 pNL4-3 based CXCR4-tropic GFP reporter virus HIV_NLGI_ and CCR5-tropic GFP reporter virus HIV_JRFL_ (kind gifts from Mary E. Klotman, Duke University) have been described previously^89^. Viral stocks were generated by transfecting HIV_NLGI_ and HIV_JRFL_ plasmids in HEK 293T cells using JetPrime transfection reagent (Polyplus). Two days later, the culture supernatant containing virus was harvested, filtered, and concentrated using a 20% sucrose cushion and ultracentrifugation at 23,000 rpm for 2 hr. The viral stock was then resuspended in PBS, aliquoted, and stored at -80° C. Viral titers were determined using a commercial p24 ELISA kit (PerkinElmer), according to the manufacturer’s instructions.

Subtype B (TRO.11), C (HIV_25710-2.43), and CRF07 (BJOX002000.03.2)^90^ HIV-1 envelope pseudoviruses were produced as previously described^91^. Briefly, 4 μg of each of the 3 HIV-envelope expression plasmids were co-transfected with 8 μg of backbone plasmid pSG3ΔEnv in HEK 293T cells using FuGene 6 transfection reagent (Promega). Virus was harvested 48 h post-transfection, filtered, and stored at −80°C with 20% FBS in the freezing media. Pseudovirus titers were determined by infecting TZM-bl-luciferase reporter cell lines and estimating 50% tissue culture infectious dose (TCID_50_) using the Bright-Glo luminescence reporter gene assay system (Promega)^92^.

To produce GFP-tagged HCMV-AD169r-BAC human cytomegalovirus (HCMV) strain, GFP-tagged AD169r HCMV BAC clone (AD169r-GFP BAC) and the CMV pp71 plasmids (kind gifts from Dr. Eain Murphy, Upstate Medical University) were co-transfected in HFF-1 cells using the Lipofectamine 3000 Transfection Reagent (ThermoFisher)^93,94^, according to the manufacturer’s instructions. Upon development of virus-induced cytopathic effect and GFP expression (∼7–10 days post-transfection), the virus was harvested, propagated twice in ARPE-19 cells, and concentrated using ultracentrifugation at 50,000 *g* for 90 minutes at 4 °C. For determining HCMV stock titers, ARPE-19 cells were seeded in 96 wells plates (20,000 cells/well) and incubated with the virus for 7 days, after which viral titers were estimated using an HCMV neutralization assay as described previously^95^.

### HIV infection of TZM-bl cells

TZM-bl-luciferase reporter cells were seeded in a 96-well plate (10^4^ cells/well) and incubated for 24 hrs. To evaluate the cytotoxicity of heat-killed bacteria, TZM-bl cells were treated every 24 hours with the equivalent of 10^4^–10^6^ CFU of unfractionated heat-killed bacteria. After 3 days, cellular luminescence was quantified using the CellTiter-Glo One Solution Assay (Promega). Cell viability was calculated by dividing the median background-corrected luminescence values by that of the negative control, with values expressed as a percentage. Cell toxicity was calculated as 100 minus the percent cell viability.

To evaluate the efficacy of heat-killed bacteria and their fractions in suppressing HIV replication, TZM-bl cells were treated every 24 hours with the equivalent of ∼10^6^ CFU of unfractionated heat-killed bacteria, washed bacterial cell pellet, or culture supernatant. After 2 days, cells were infected with 5 ng of p24-equivalent HIV_NLGI_ or HIV_JRFL_. Cells treated with an ARV cocktail (10 μg/μL zidovudine, 10 μg/μL lamivudine, and 1 μg/μL nevirapine; Sigma) were used as a positive control. Two days after HIV infection, viral replication was quantified using the luciferase reporter assay described above.

To evaluate the efficacy of heat-killed bacteria in suppressing replication of different HIV clades, TZM-bl cells were infected with TRO.11 (TCID_50_: 2223.05), HIV_25710-2.43 (TCID_50_: 2344.05), or BJOX002000.03.2 (TCID_50_: 3493.85). Cells were also treated with 10^6^ CFU-equivalent of heat-killed bacteria or ARV cocktail. After 26 hours, viral infection was quantified using the luciferase reporter assay described above.

To evaluate the efficacy of isogenic bacterial mutants and purified small molecules in modulating HIV replication, TZM-bl cells were incubated with HIV_JRFL_ (5ng of p24-equivalent) and 10^6^ CFU-equivalent of heat-killed bacteria or small molecules (1 nM of DL-indole-3-lactic acid [ILA], 1 nM of indole-3-carboxaldehyde [I3A], 1 nM of 6-formylindolo[3,2-b]carbazole [FICZ], or 10 μM of CH223191; Sigma). After 26 hours, viral infection was quantified using the luciferase reporter assay described above.

In each independent assay, every treatment has 3–6 technical replicates. HIV replication was calculated by dividing the median background-corrected luminescence values by that of the negative control, with values expressed as a percentage. Percent inhibition of HIV replication was calculated as 100 minus the percent HIV replication.

### Flow cytometric analysis of HIV-infected TMCs treated with bacteria

TMCs were resuspended in RPMI 1640 (Gibco), supplemented with 10% FBS, antibiotic/antimycotic mixture (Gibco), 1 mM of sodium pyruvate (Sigma-Aldrich), and non-essential amino acids (Gibco). TMCs were seeded into a flat-bottom 96-well plate (10^6^ TMCs/well), infected with 250 ng of p24-equivalent of GFP-tagged HIV_NLGI_, and treated with either 10^6^ CFU-equivalent of heat-killed bacteria or the ARV cocktail. Plates were then centrifuged at 1200 *g* for 2 hr to help synchronize infection. After 48 hours, TMCs were harvested, incubated with anti-human CD4 (BioLegend) at 4°C for 30 min, washed twice with PBS, stained with LIVE/DEAD Fixable Aqua Dead Cell Stain Kit (ThermoFisher) according to the manufacturer’s instructions, and fixed using 10% formalin. The stained TMCs were analyzed on an LSRFortessa (BD Biosciences) using BD FACS Diva software. Data were analyzed with FlowJo software (version 10.8.1; Tree Star, Inc).

### Time-of-addition assay

TZM-bl-luciferase reporter cells were seeded in a 96-well plate (10^4^ cells/well) and incubated for 24 hrs, after which they were incubated with 5 ng of p24-equivalent HIV. After 1 hr, the virus-containing supernatant was removed, cells were washed twice with Dulbecco’s PBS (DPBS), and fresh media was added. HIV-infected cells were then treated with anti-HIV drugs (1 μM of enfuvirtide, 1 μM of zidovudine, or 1 μM of raltegravir; MedChemExpress), 2.5 μg of the HIV gp120 CD4-binding site-specific monoclonal antibody VRC01 (BEI Resources), or 10^6^ CFU-equivalent of heat-killed bacteria at 0, 2, 4, 6, 8, 10, or 12 hours. The degree of viral infection was quantitated at 26 hours post-HIV infection using the luciferase reporter assay described above.

In each independent assay, each treatment has 3 technical replicates. HIV replication at a certain time-point post-infection was calculated by dividing the median background-corrected luminescence values by the same treatment added at 12 hours post-infection, with values expressed as a percentage. Percent inhibition of HIV replication was calculated as 100 minus the percent HIV replication.

### Generation of *C. immunis*ΔArAT

We used homologous recombination to disrupt the ArAT gene in *C. immunis*, using pMTL82151, one of the ClosTron plasmids^96^, as a suicide vector. We used the Q5 High-Fidelity DNA polymerase (NEB) to PCR amplify the *ermB* gene that provides resistance to erythromycin (using pMTL83251 as the template) and two 800 bp regions (separated by 80 bp) that flanked the *C. immunis* ArAT target site (representing left and right homology arms), using genomic DNA from *C. immunis* as the template. PCR primers used are as follows: ArAT Left F: gctcggtacccggggatcctCTCTGACTGTATTCTATGAC; ArAT Left R: cttcggccggGTCTTTCCGTTTATTTTCTC; ArAT Right F: gaatgtgtttTCCTGGAAAAGAAGGAAAAAG; ArAT Right R: agcttgcatgtctgcaggccCGGCATGAAAAATCCCAG; ErmB F: acggaaagacCCGGCCGAAGCAAACTTAAG; ErmB R: ttttccaggaAAACACATTCCCTTTAGTAACGTG These three PCR products were digested with XbaI and XhoI and cloned into XbaI- and XhoI-digested pMTL82151 plasmid using HiFi assembly (NEB). Electrocompetent *E. coli* S17λPir was transformed with the ligation mixture, which generated S17λPir:pMTL82151ΔArAT. This strain was used to transfer pMTL82151ΔArAT into *C. immunis* using standard conjugation methods. Briefly, overnight cultures of *E. coli* donors and *C. immunis* recipients were mixed at a 10:1 donor-to-recipient ratio. Multiple aliquots (30 μl) of the mixture were spotted onto a BHI-S agar plate and incubated anaerobically at 37°C. After 24–48 h, the total growth on each plate was collected, resuspended in fresh media, and grown on BHI-S plates containing erythromycin (500 μg/ml) and colistin (10 μg/ml). To confirm that the plasmid was not present in *C. immunis*, we ensured the culture did not grow on BHI-S plates containing colistin (10 μg/ml) and chloramphenicol (12.5 μg/ml), against which the plasmid backbone confers resistance. Erythromycin-resistant colonies of *C. immunis* were confirmed to harbor the insertion of the *ermB* gene and expected 80-bp deletion in ArAT by PCR and Sanger sequencing.

### Metabolomics of bacterial culture supernatants

To validate the loss-of-function of *C. immunis*ΔArAT, overnight cultures of wildtype *C. immunis* and *C. immunis*ΔArAT were grown in PYG and pelleted to obtain cell-free supernatants. Untargeted metabolomics of the respective supernatants was performed via gas chromatography/mass spectrometry (GC/MS) at the Duke Molecular Physiology Institute. Proteins in aliquots of 100 µL culture supernatant in surface-inactivated glass GC vials (MicroSolv, Leland, NC) were crash-precipitated by addition of 750 µL of dry methanol spiked to 6.25 mg/L perdeuterated myristate as a retention-time-locking (RTL) standard. After centrifugation at 13,500 *g* for 10 minutes (MicroSolv Vial Centrifuge, Leland, NC), clear extracts were transferred to fresh glass GC vials and vacuum-dried in a SpeedVac (ThermoFisher), followed by azeotropic drying with 150 µL toluene per sample. Dry extracts were methoximated, trimethylsilylated, analyzed in GC/MS, and the data reduced, as previously described^97,98^. Identity of the key metabolite, indole-3-lactic acid (ILA), was confirmed by comparison to an authentic standard by its *tris*-trimethylsilyl derivative’s retention time of ∼20.077 minutes, combined with its its molecular ion at *m/z* 421, with characteristic heavy fragments at *m/z* 202 and 304.

### *Cdk1* and *Cdk2* expression analysis

TZM-bl luciferase reporter cells (2 x 10^5^ cells/well) were incubated with ∼10^6^ CFU-equivalent of heat-killed bacteria and infected with 5 ng of p24-equivalent HIV_NLGI_ virus. After 24 hr, RNA was extracted using an RNeasy mini kit (Qiagen). cDNA was prepared using the SuperScript™ III First-Strand Synthesis System (ThermoFisher), and qPCR was performed on a QuantStudio 7 Flex real time PCR machine (ThermoFisher) using TaqMan gene expression mastermix (ThermoFisher) and TaqMan Gene Expression Assays (Applied Biosystems) for *Gapdh*, *Cdk1*, and *Cdk2*. The thermal cycling steps were 50°C for 2 min, 95°C for 10 min, and 40 cycles of 95°C for 15 s and 60°C for 1 min. The comparative Ct method was used to quantify transcripts that were normalized with respect to *Gapdh*.

### Bacterial treatment of HCMV-infected ARPE-19 cells

The efficacy of *C. immunis* and *C. immunis*ΔArAT to modulate human cytomegalovirus (HCMV) replication was assessed by performing the HCMV neutralization assay as previously described^95^. Briefly, ARPE-19 cells (5000 cells/well) were seeded in 384-well black plates with clear bottoms (Corning) and cultured for 24 hours. ARPE-19 cell monolayers were then incubated with GFP-tagged HCMV-AD169r-BAC virus strain at a multiplicity of infection (MOI) of 1.5 along with either ∼10^6^ CFU-equivalent of heat-killed bacteria or 0.625 mg/mL of anti-CMV immune globulin (Cytogam; CSL Behring Healthcare). After 48 hrs, cell culture media was aspirated and cells were fixed with 10% formalin at room temperature for 15 min. Fixed cells were washed twice with PBS + 0.1% Tween 20 and incubated with 1:1000 mouse anti-HCMV IE-1 monoclonal antibody (clone 8B1.2; Millipore) in staining buffer (DPBS containing 1% FBS and 0.3% Triton X-100) for 1 hr at RT. After subsequent washing, cells were stained with goat anti-mouse IgG-AF488 (Abcam) at 1:1000 dilution for 1 hr at RT. After further washing, the cell nucleus was stained with 1:1000 4’,6’-diamidino-2-phenylindole (DAPI; Invitrogen) for 10 minutes at RT. DAPI-positive (indicative of total cells) and AF488-positive cell counts (indicative of HCMV-infected cells) were measured using ImageXpress Pico Automated Cell Imaging System (Molecular Devices), and the percentage of HCMV-positive cells of total cells was computed. In each independent assay, every treatment has ≥2 technical replicates. CMV replication was calculated by dividing the median background-corrected values by that of the negative control (i.e., cells treated with *B. fragilis*) with values expressed as a percentage. Percent inhibition of CMV replication was calculated as 100 minus the percent CMV replication.

### Analysis of ArAT abundances in metagenomic datasets

For individuals at-risk for HIV, we downloaded a publicly available shotgun metagenomic dataset from the mSTUDY cohort that longitudinally followed men who have sex with men for the acquisition of HIV^40^. For individuals who did not acquire HIV (control), we analyzed the first sample available. For men who acquired HIV (cases), we analyzed samples from the visit immediately preceding seroconversion.

For children undergoing hematopoietic stem cell transplantation (HSCT), the longitudinal metagenomic dataset has been previously described^43^. Briefly, we obtained weekly fecal samples beginning up to one month before transplant and continuing until 100 days post-transplant. Shotgun metagenomic data were previously generated^43^. Patients were assessed for CMV viremia by weekly PCR testing of blood. We excluded patients who had detectable CMV ≥5 days prior to transplant or ≥28 days following transplant. Informed consent was obtained from participants’ legal guardians prior to enrollment, and the study protocol was approved by the DUHS IRB (Pro00064365).

For individuals with household exposure to SARS-CoV-2, we utilized fecal samples collected from a prospective cohort study previously described^45^. In brief, participants (age range, 0–58 years old; median, 10.5 years old) were identified through review of the results of SARS-CoV-2 testing conducted within the DUHS from April 2020 to June 2022. All individuals identified in the household who had close contact with a SARS-CoV-2–infected person were eligible for study enrollment. Participants completed questionnaires regarding SARS-CoV-2 exposures and symptoms. Stool samples were collected into 100% ethanol and RNAProtect (Qiagen) using EasySampler Stool Collection Kits (ALPCO Diagnostics). Individuals who were positive for SARS-CoV-2 by PCR testing of nasopharyngeal swabs and had self- or caregiver-reported fever, cough, shortness of breath, sore throat, rhinorrhea, nasal congestion, headache, abdominal pain, vomiting, diarrhea, anosmia, dysgeusia, chest pain, and/or myalgias were considered symptomatic. Individuals who tested positive for SARS-CoV-2 but did not have any of these symptoms were considered to have asymptomatic infection. We focused on samples collected from participants who had not taken antibiotics in the 30 days prior to study enrollment. The study was approved by the DUHS IRB (Pro00105249, Pro00106150).

For the latter two studies, DNA was extracted from the fecal samples using PowerSoil Pro Kits (Qiagen). Shotgun metagenomic sequencing libraries were constructed using Nextera XT DNA Library Prep Kits (Illumina) and sequenced on NextSeq500 or NovaSeq6000 instruments (Illumina) as 150-bp paired-end reads. For all three datasets, we trimmed sequence reads and removed host decontamination using KneadData. Forward and reverse reads that passed quality control (paired and unpaired) were concatenated, and samples with <25,000 reads were discarded. The resulting reads were fed into the HUMAnN 3 pipeline for functional profiling as UniRef90 gene families^99^. To identify homologs of the ArAT gene, we used the *C. immunis* and *L. reuteri* ArAT gene sequences to search for UniRef90 entries in the UniProt Knowledgebase^100^, setting a similarity threshold of ≥30%. We used the grep UNIX command to extract the abundance of these ArAT homologs from the library-depth normalized UniRef90 gene family abundance table, and we summed their abundances. Any homologs identified by both the *C. immunis* and *L. reuteri* ArAT gene sequence were only counted once.

For the longitudinal dataset from children undergoing HSCT, we used the gam function of the mgcv R package to fit a generalized additive mixed model (gamm), with the total ArAT abundance as the response. For the predictors, we used a fixed effect for group, a smooth term for days relative to transplant to model the longitudinal data, and a random effect for each individual to account for within-person correlation. The difference in ArAT gene abundance between the CMV-infected and CMV-uninfected groups was assessed by a Wald test of the fixed effect group coefficient. Results were graphed using the ggplot2 R package.

### Statistical Analysis

The specific statistical tests used are detailed in each figure legend. Prism 10 (GraphPad Software) was used for all statistical analyses, with the exception of the gamm used for the pediatric HSCT metagenomic data and the viral replication assays (HIV and CMV), which were both done in R (version 4.2.2). For the latter, we performed paired t-tests (by individual experiment) on the background-corrected luminescence values, comparing the intervention to its relevant negative control; we applied the Benjamini-Hochberg procedure for multiple comparison correction. We consider *P*>0.1 as not significant. *P*-values between 0.05 and 0.1 are indicated explicitly on each figure.

### Data Availability

The 16S rRNA gene sequencing data for the SHIV-challenged infant rhesus macaques are available in the NCBI Sequence Read Archive (SRA) repository, accession number PRJNA1027904; 16S rRNA gene sequencing data for the Kenyan children cohort are deposited in the NCBI SRA repository, accession number xxxxxx. The shotgun metagenomic sequencing data from the HSCT and COVID-19 cohorts presented in the study are deposited in the NCBI SRA repository, accession numbers PRJNA890666 and PRJNA948840, respectively. The data from the mStudy cohort was previously published^40^; the metagenomic data were downloaded from BioProject PRJNA836336, and the corresponding metabolomic data were downloaded from Dryad (https://doi.org/10.5061/dryad.np5hqbzx5).

## ACKNOWLEDGEMENTS

This work was supported by Gilead Sciences Research Scholar’s Program in HIV (RG), National Institutes of Health (NIH) grants P01 AI117915 (KVR, MGH, KDP, SRP) and P01 AI178377 (KVR, MGH, KDP, SRP, RG, NKS), a pilot award from the Duke Center for AIDS Research (CFAR; P30 AI064518) and Duke Microbiome Center (NKS), and The Office of Research Infrastructure Programs/OD grant P51OD011107 to the CNPRC. DJ received funding from the Duke CFAR Summer Internship program (R25 AI140495). CYT received support from American Heart Association pre-doctoral fellowship AHA 897275, a National University of Singapore Development Grant, and a Tan Kah Kee Postgraduate Scholarship. JRB received support from NIH 5P30DK124723, 5R01DK117491, 1U24DK129557, and 2P30AG027816, and USDA 2020-28640-31521. The funders had no role in study design, data collection and interpretation, or the decision to submit the work for publication; the content is solely the responsibility of the authors. We thank the Duke Microbiome Core Facility for performing the DNA extractions, quality control, and preparing the 16S rRNA gene sequencing library; the Duke Sequencing and Genomic Technologies Shared Resource for sequencing the 16S rRNA gene library as well as preparing and sequencing the shotgun metagenomic libraries; and the Duke Compute Cluster for providing the high-performance computing hardware used in the bioinformatic analysis of metagenomic sequencing data. The following reagents were obtained through BEI Resources, NIAID, NIH: TZM-bl cells (ARP-8129, contributed by Dr. John C. Kappes, Dr. Xiaoyun Wu and Tranzyme Inc.) and monoclonal anti-HIV-1 gp120 protein (VRC01, produced in vitro; ARP-12033).

## CDC Disclaimer

This work was supported in part by the Centers for Disease Control and Prevention (CDC). The findings and conclusions in this report are those of the author and do not necessarily represent the official position of the CDC.

## AUTHORS CONTRIBUTIONS

Conceptualization: D.J., R.G. and N.K.S.

Data curation: D.J.

Formal analysis: D.J., K.X., M.G.H.

Funding acquisition: D.J., K.V.R., K.D.P., and S.R.P., R.G., and N.K.S.

Investigation: D.J., N.S., C.Y.T., S.D., H.Y.W., B.S.T., D.H., J.R.B., and R.G.

Resources: NHP fecal samples: A.A., K.V.R., K.D.P., and S.R.P.; fecal samples and clinical metadata from Kenyan children: J.P.S. and R.S.; metagenomic and clinical data for pediatric HSCT patients: S.M.H. and M.S.K.; metagenomic and clinical data for SARS-CoV-2-exposed individuals: D.J., J.H.H., J.R., M.S.K, and N.K.S.; *L. gasseri* strains: N.Y.S.; *L. reuteri* strains: L.C.-B.

Supervision: R.G. and N.K.S.

Writing – original draft: D.J., N.S., C.Y.T., R.G. and N.K.S.

Writing – review & editing: all authors.

## COMPETING INTERESTS

MSK serves as a consultant for Merck. SRP serves as a consultant for Moderna, Pfizer, Dynavax, and GSK on their CMV vaccine programs, and has led sponsored research programs with Moderna, Dynavax, and Pfizer on CMV vaccines. DJ, CYT, SRP, RG, and NKS are inventors on a patent application submitted by Duke University that covers the therapeutic use of ArAT-expressing bacteria and their metabolites in viral infections. Other authors have no conflict of interest to disclose.

## Supplementary Figure Legends

**Supplementary Figure 1. C. immunis and R. gnavus inhibit replication of different HIV clades without inducing cytotoxicity, related to Figure 2. (A)** Percent cytotoxicity in TZM-bl cells treated with the indicated commensal bacteria at various multiplicities of infection (MOI). The dashed line indicates 25% cytotoxicity. **(B–D)** Percent inhibition of HIV replication in TZM-bl cells treated with the indicated commensal bacteria. HIV strains used are from clade B **(A)**, clade CRF07 **(B)**, and clade C **(C)**. Data are normalized to cells treated with *B. fragilis*. ARV, antiretroviral cocktail. Data in panels B–D are from 6 independent experiments, with bars representing means ± SEM. *, *P*<0.05; **, *P*<0.01 by paired t-tests for each treatment compared to the negative control and adjusted by the Benjamini-Hochberg procedure.

**Supplementary Figure 2. The kinetics of the HIV replication cycle as defined by canonical anti-HIV drugs, related to Figures 2 and 3.** Time-of-addition assay results for VRC01 (an anti-CD4 antibody), enfuvirtide, zidovudine, and raltegravir. Data are normalized to cells treated with the same drug at 12 hours, are from 3 independent experiments, and are represented as means ± SEM. RT, reverse transcription; IN, integration. *, *P*<0.05 by paired t-tests.

**Supplementary Figure 3. Individuals who ultimately acquire HIV have decreased fecal levels of Lachnospiraceae and serum levels of ILA, related to Figure 5. (A)** The fecal abundance of the bacterial family Lachnospiraceae in adults at-risk for HIV who did (case; *n* = 9) or did not acquire HIV (control; *n* = 20). **(B)** Serum abundance of ILA in the same adults at-risk for HIV who did or did not acquire HIV. In both panels, the samples for the cases were obtained before they tested positive for HIV. *, *P*<0.05 by Mann-Whitney U test **(A, B)**.

**Supplementary Figure 4. SARS-CoV-2 infection status does not alter the fecal abundance of the ArAT gene, related to Figure 5.** The fecal abundance of the ArAT gene in individuals who were (*n* = 233) or were not (*n* = 52) infected with SARS-CoV-2. ns, not significant by Mann-Whitney U test.

## Notes

### Summary of Updates

This version of the manuscript has been revised to include additional analysis and data.

