## Supplementary figures and images for "Commensal bacteria inhibit viral infections via a tryptophan metabolite"

### Supplemental figures

**Supplementary Figure 1**

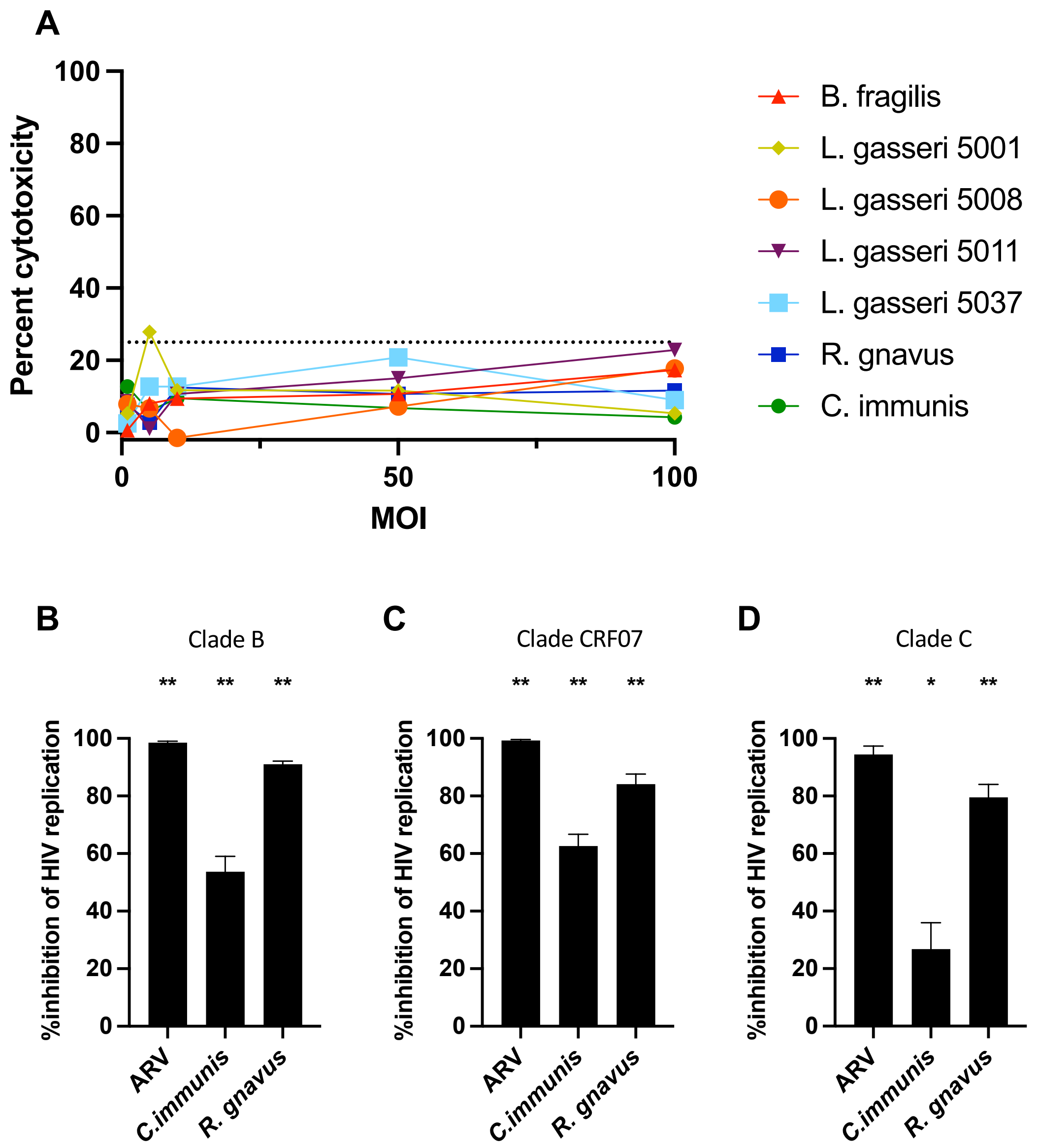

#### Supplementary Figure 2

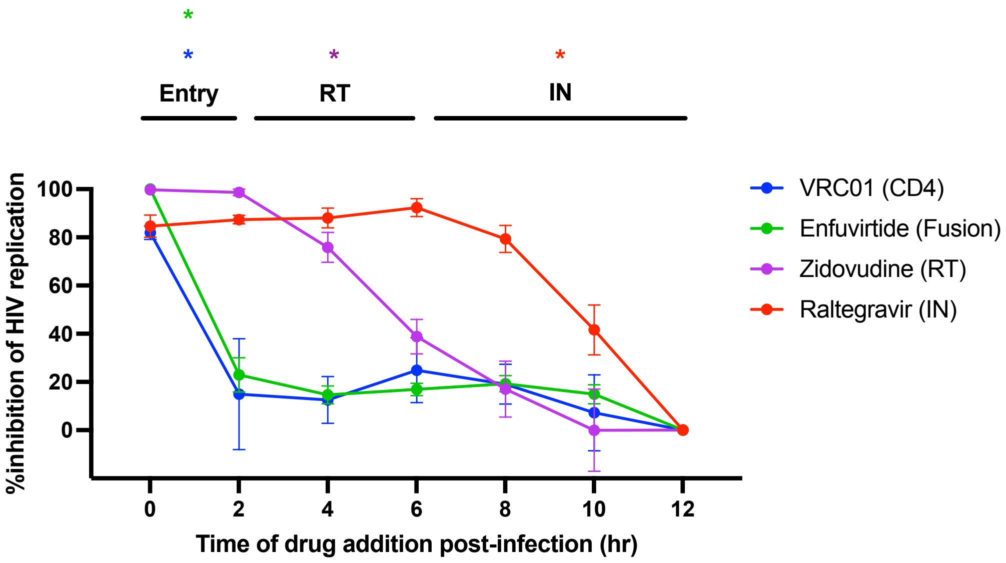

**Supplementary Figure 3**

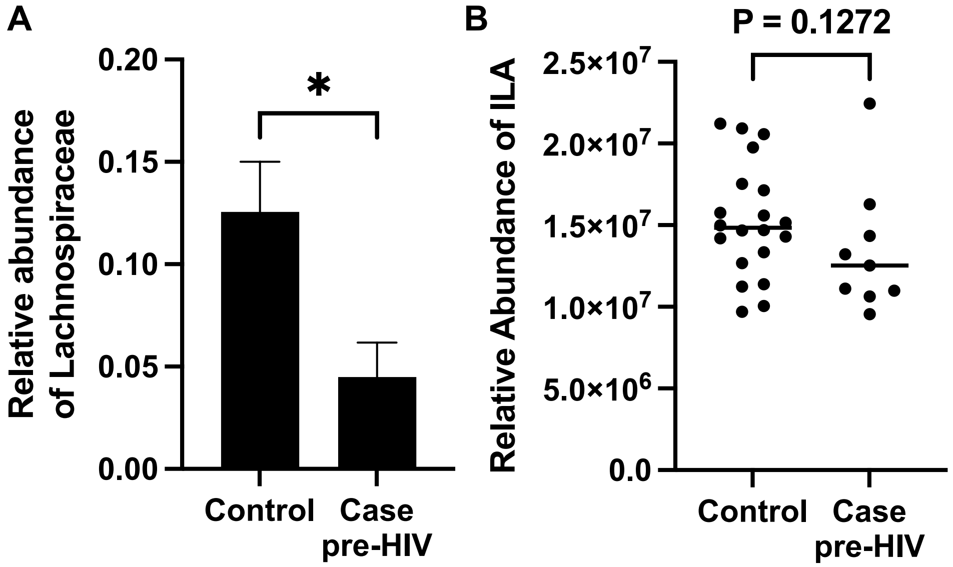

**Supplementary Figure 4**

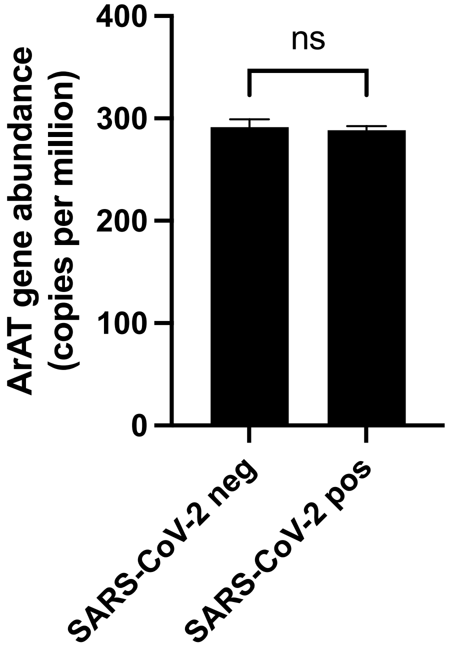
