## Supplemental table for "Commensal bacteria inhibit viral infections via a tryptophan metabolite"

**Supplementary Table 1. Differentially abundant taxa present in different pairwise comparisons used in the microbe-phenotype triangulation approach, related to Figure 1.**

| |
|---|
| **Vax early-infected (*n*=6) vs. Vax late-infected (*n*=4)** |
| 4 taxa more abundant in late-infected animals |
| *Lactobacillus gasseri* |
| Lachnospiraceae |
| *Bacteroides vulgatus* |
| *Bacteroides sp.* |
| **Ctl early-infected (*n*=7) vs. Ctl late-infected (*n*=1)** |
| n/a |
| **Vax early-infected (*n*=6) vs. Ctl early-infected (*n*=7)** |
| 6 taxa differentially abundant between the two groups |
| *Clostridium sensu stricto* |
| *Prevotella sp.* |
| *Rikenellaceae* RC9 gut group |
| *Bacteroides vulgatus* |
| *Sutterella sp.* |
| *Bacteroides sp.* |
